# Cellular Population Dynamics Control the Robustness of the Stem Cell Niche

**DOI:** 10.1101/021881

**Authors:** Adam L. MacLean, Paul Kirk, Michael P.H. Stumpf

## Abstract

Within populations of cells, fate decisions are controlled by an indeterminate combination of cell-intrinsic and cell-extrinsic factors. In the case of stem cells, the stem cell niche is believed to maintain “stemness” through communication and interactions between the stem cells and one or more other cell-types that contribute to the niche conditions. To investigate the robustness of cell fate decisions in the stem cell hierarchy and the role that the niche plays, we introduce simple mathematical models of stem and progenitor cells, their progeny and their interplay in the niche. These models capture the fundamental processes of proliferation and differentiation and allow us to consider alternative possibilities regarding how niche-mediated signalling feedback regulates the niche dynamics. Generalised stability analysis of these stem cell niche systems enables us to describe the stability properties of each model. We find that although the number of feasible states depends on the model, their probabilities of stability in general do not: stem cell–niche models are stable across a wide range of parameters. We demonstrate that niche-mediated feedback increases the number of stable steady states, and show how distinct cell states have distinct branching characteristics. The ecological feedback and interactions mediated by the stem cell niche thus lend (surprisingly) high levels of robustness to the stem and progenitor cell population dynamics. Furthermore, cell–cell interactions are sufficient for populations of stem cells and their progeny to achieve stability and maintain homeostasis. We show that the robustness of the niche — and hence of the stem cell pool in the niche — depends only weakly, if at all, on the complexity of the niche make-up: simple as well as complicated niche systems are capable of supporting robust and stable stem cell dynamics.

## Introduction

Stem cells control the essential processes that facilitate multi-cellular life. Their ability to continue to produce more specialised types of cells in a coordinated manner underlies developmental processes, tissue regeneration and wound repair. Stem cell function relies crucially on the ability to make robust cell fate choices (Gurtner et al., 2008; Reya et al., 2001). These include the choice between self renewal and differentiation, or — once committed to differentiation — the choice between two or more specialised cell lineages. Many factors compound the decision-making process, ranging from cell-intrinsic regulation, to cell-extrinsic factors such as intercellular signalling and environmental stresses (Enver et al., 2009). Failure to make such cell fate choices correctly, by contrast, leads to disease and interferes with a host of physiological processes, ranging from control of the immune response to normal and healthy ageing (Brack et al., 2007; Geiger et al., 2013; Uccelli et al., 2008).

Stem cell function is therefore safeguarded by a number of mechanisms, that include an apparently delicately balanced interplay with other cells. The concept of the stem cell niche (Schofield, 1978), has proved vital to our understanding of stem cell function and maintenance in a variety of cycling tissues including blood, skin and intestine (Reya et al., 2001; Spradling et al., 2001). Indeed, it can be argued that the ability of a cell to exhibit *stemness* cannot be defined in isolation — that is, without considering the influence of the niche. Conceptually, niches can be treated as domains of influence in which different cell populations can reside and exert effects on one another though signalling, paracrinal and juxtacrinal interactions. Using such a description of stem cell niches, we can test hypotheses regarding their particular extent, form, and constituents. For example, we can now test if the interactions that maintain stemness (or the niche) are indeed carefully balanced, or show a level of robustness to external perturbations. Given the overlap between niches as defined in stem cell biology and population biology (Székely et al., 2014), the population biological viewpoint lends itself well to analysis of stem cell systems.

Population biology has a rich history of applications to a wide range of systems from ecological networks to social organisations (May, 1972; MacArthur & Wilson, 1967; MacLean et al., 2013; Nowell, 1976; Saavedra et al., 2011); the question whether such systems are robust or fragile has been central to may of these studies. Populations — whether they are composed of animal species or cell species — obey certain principles that both determine and are affected by the birth, growth, and death characteristics. These may be complex functions that depend on interactions with other species, which can be either positive (mutualistic), negative (competitive), or a mixture of the two. By integrating these processes we can build up a description of the dynamics of interacting populations. Stem cells and their progeny are well-suited to such a description and population biological concepts begin to gain a foothold in stem cell biology (MacLean et al., 2013; Mangel & Bonsall, 2013).

The stability of a population dynamical system refers to its ability to return to the same state following some small perturbation away from its point of equilibrium (Strogatz, 1994). This is an important concept: it enables us to measure the robustness of a particular state, and to investigate which states may be capable of persisting in nature. There has been much debate over whether increasing the complexity of a system is likely to lead to an increase in stability. Early work supported this hypothesis (Elton, 1958; MacArthur, 1955); May subsequently proved a theorem stating that, in general, the stability of a system will decrease as its complexity increases (May, 1972). More recent results have extended May’s result and suggested that the system stability can change dramatically when specific types of interaction are considered (Allesina & Tang, 2012; Kirk et al., 2015, in press). In particular, Kirk et al. (Kirk et al., 2015, in press) relax the assumption that species only interact with one another at random, and in doing so move closer to a description of systems that we might expect to find in the natural world.

In order to investigate the stability of stem cell states, we develop models describing the dynamics of a stem cell lineage and study their equilibrium states – which vary between models – in order to determine which states can persist in nature. Since the stability of each state depends upon the values taken by the model’s parameters, it is necessary to consider a range of biologically feasible parameter values. The *stability probability* for a given state is defined to be the proportion of times that the state was found to be stable after repeatedly sampling parameter values from within this range (Kirk et al., 2015, in press; Christianou & Kokkoris, 2008; Gilpin, 1975; Pimm, 1984; Roberts, 1974).

The crucial stem cell processes of self renewal, differentiation and lineage choice are of particular interest in the models developed here. We introduce four population biological models that share these characteristics but differ in their number of lineages and feedback characteristics. Structure within these models is identified as a key factor in maintaining stability. Further analysis of one of the models demonstrates how different stable states can be reached from different experimental conditions (corresponding to different parameter values). This provides insight into how stem cells maintain homeostasis and how multiple states can be accessed, and could explain how, for example, depletion of a particular blood cell population is remedied at the stem/progenitor cell level by a state shift to one that repopulates the haematopoietic system.

## Results

### Fixed points in the stem cell hierarchy define stable cell states

In order to assess the stability of cell states, we study the fixed points of model systems. Fixed points correspond to invariant states that are reached as a system approaches stationarity (other stationary states – such as oscillations or *limit cycles* – are also possible). In Figure 1 we give an illustration of fixed points: these are the minima of the state space defined by a potential function, and cells lying within a basin of attraction will evolve towards them. Lower minima may correspond to terminally differentiated cell states, and higher minima — with higher potential — to stem or progenitor cell states. They can also be thought of as the local (or global) minima in Waddington’s landscapes (Waddington, 1957) — but here the landscape corresponds to the population dynamics, and not the intra-cellular dynamics of stem cell behaviour as characterised by (e.g.) stem cell markers Nanog and Pecam for embryonic stem cells, or CD34 and Sca1 for haematopoietic stem cells (Rué & Martinez-Arias, 2015).

**Figure 1:**
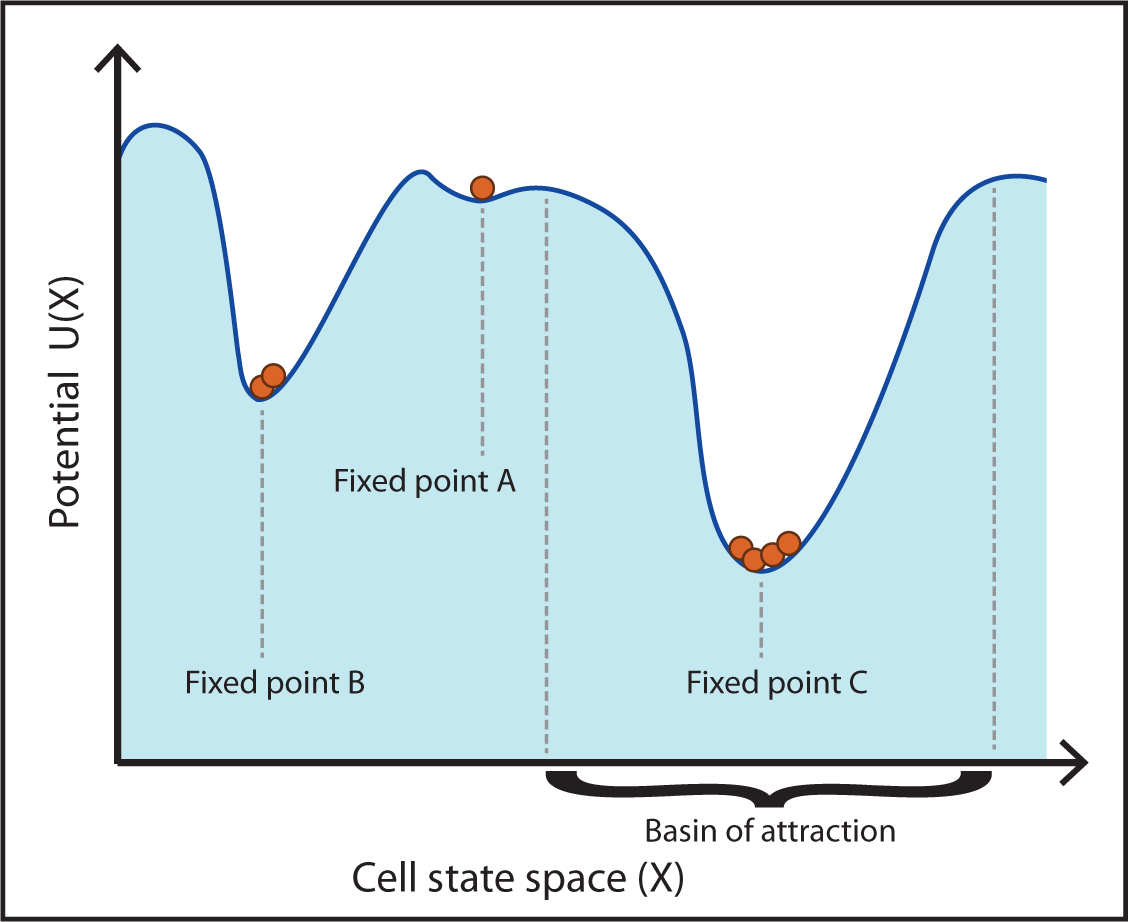
Fixed points in cell state space. Fixed points occur at minima on the landscape of cellular states, and correspond to persistent phenotypes. Each fixed point has a basin of attraction that defines the extent of its reach. Here, fixed point A may correspond to a stem or progenitor cell state, and fixed points B and C (with lower energy minima) to terminally differentiated cell states.

We consider typical, albeit simplified, stem cell differentiation hierarchies, consisting of three cell populations: stem cells (*S*), progenitor cells (*P*), and differentiated cells (*D*). Four models are constructed from these cell populations, shown in Figure 2; these differ in their branching and feedback characteristics (Buzi et al., 2015). The models are, of course, (vastly) simplified descriptions of more complicated processes, however they serve our goal to compare characteristics, and as such can provide insight into basic mechanisms of stem cell function. The details of and assumptions underlying the models are given in the Methods. For the analysis of fixed points of a system, we have developed methods of generalised stability analysis that allow us to characterise the fixed points of stem cell models and assess their stability (Kirk et al., 2015, in press). We provide a description of these methods and the statistical procedures that we use in Methods. The crucial concept derived from these methods is the stability probability of a fixed point. This defines the probability that a fixed point of a model will be stable, given that we know that the model parameters will lie within some range, but we do not know their values.

**Figure 2:**
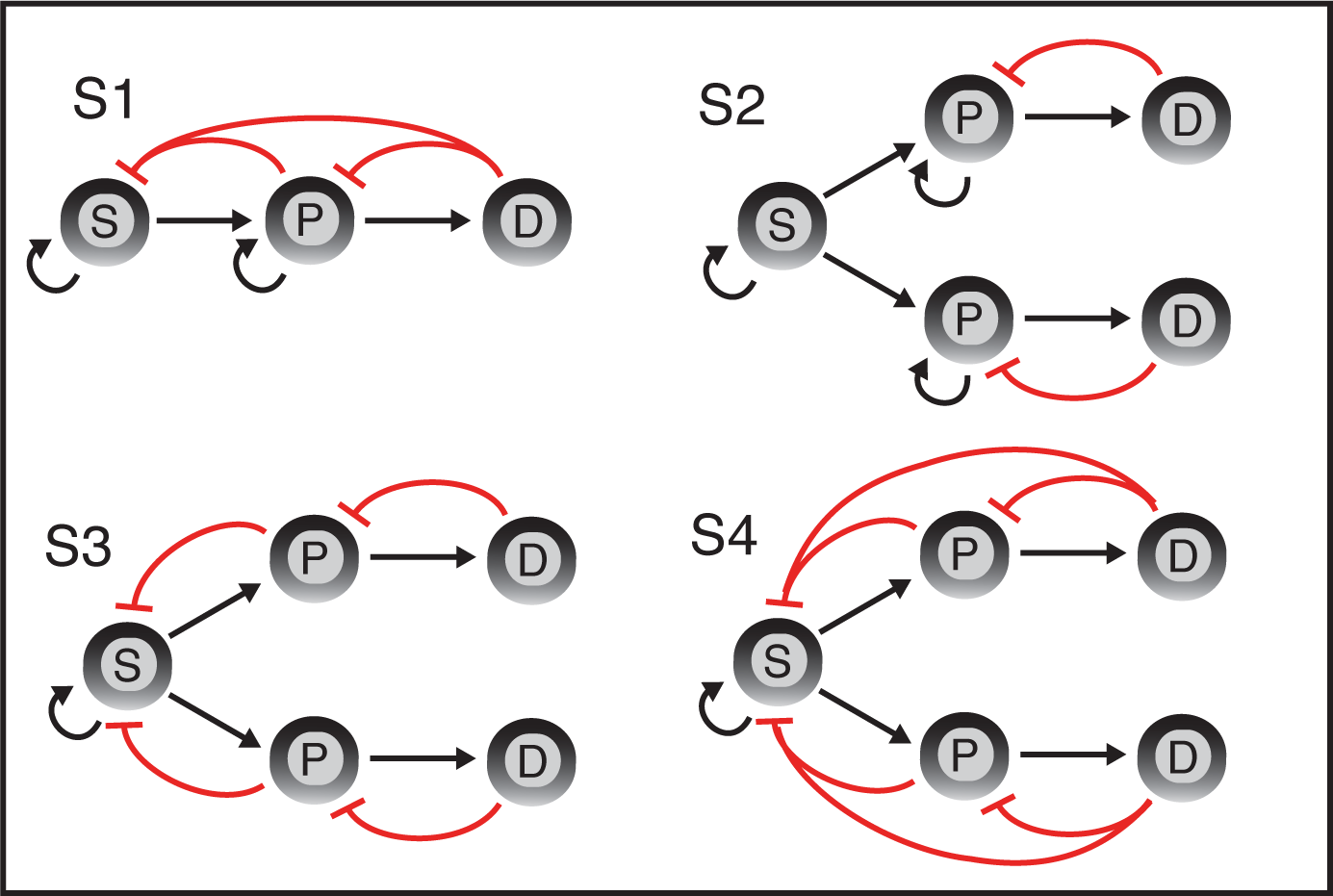
*Schematic description of the models S*1 – *S*4. Black arrows denote production by proliferation or differentiation and red arrows denote inhibition of cell proliferation by negative feedback.

Shown in Table are the stability probabilities for each fixed point of the models under investigation in column *a*. The number of fixed points differs between models *S*1 *– S*4. Each model has the origin as a fixed point (fixed point 1); this corresponds to a state where all species go extinct. We do not analyse these points further since they are not of biological interest (some might still have interesting mathematical properties). In addition fixed point 2 for models *S*3 and *S*4 is not reachable; that is, the system will never end up in this state starting within the parameter ranges that we study. In previous work, Roberts (Roberts, 1974) referred to reachable fixed points as *feasible*. Here we will also leave aside the unreachable fixed points, and proceed to analyse those fixed points that are both reachable and nonzero.

The number of biologically meaningful (nonzero and reachable) cell states is two for models *S*1, *S*3, and *S*4, and one for model *S*2. We see that the number of cell types modelled does not correspond to the number of possible fixed points. For model *S*1 (see Figure 2), fixed point 2 describes a state where progenitor and differentiated cell populations co-exist, but the stem cell pool has become completely depleted. The probability that this fixed point is stable is 0.75. The second fixed point describes a state where all three cell populations are positive, and this state will always be stable no matter where one begins in parameter space.

Of the three models that each represent five populations, model *S*2 has only one relevant fixed point — even fewer than model *S*1. This tells us that interactions only between differentiated and progenitor cells and not with the stem cell compartment limits the richness of dynamics available. The single biologically relevant state of model *S*2 is always stable. Model *S*3 has two reachable fixed points, both of which have positive population sizes for all five species (we now have branching in the stem cell compartment into two progenitor cell species). Each of these fixed points is stable for all parameter values: the system, by virtue of the nature of the cell–cell interactions is robust. For the final model, *S*4, we see that its stability properties closely reflect those of model *S*3: both biological fixed points (3 and 4) are always stable. Thus, the presence or absence of a direct signal from differentiated cells onto stem cells does not have great effect on cell state stability.

### Structure in the stem cell hierarchy maintains stability

In addition to the true stability probabilities obtained for each model state (Table, column *a*), we also calculate the stability probabilities under models that ignore the statistical dependencies inherent to real dynamical systems; this perspective has been very popular in population biology, where it was often (Allesina & Tang, 2012; May, 1972) (but not always (Kirk et al., 2015, in press; Roberts, 1974)) seen as a valid attempt at assessing the stability of ecological systems. There the surprising result has been that large and complex ecological systems tend to be less stable than simple systems; the results in columns *b* and *c* in Table correspond to the stability probabilities obtained for such models. See Methods for a description of how each of these distributions was calculated.

**Table 1:** The stability probabilities for each fixed point of models *S*1 – *S*4. For each fixed point: (*a*) is the true stability probability; (*b*) is the stability probability under an independent null distribution; (c) is the stability probability under an *i.i.d*. null distribution.

| Model | Fixed Point | (a) | (b) | (c) |
| --- | --- | --- | --- | --- |
| S1 | 1 | 0.25 | 0.25 | 0.034 |
|  | 2 | 0.75 | 0.75 | 0.13 |
|  | 3 | 1.0 | 0.94 | 0.19 |
| S2 | 1 | 0.83 | 0.83 | 0.0038 |
|  | 2 | 1.0 | 1.0 | 0.011 |
| S3 | 1 | 0.83 | 0.83 | 0.0038 |
|  | 2 | — | — | — |
|  | 3 | 1.0 | 1.0 | 0.018 |
|  | 4 | 1.0 | 1.0 | 0.018 |
| S4 | 1 | 0.83 | 0.83 | 0.0036 |
|  | 2 | — | — | — |
|  | 3 | 1.0 | 0.96 | 0.023 |
|  | 4 | 1.0 | 0.96 | 0.023 |

Upon comparison of columns *a – c* in Table, two main observations can be made. First, there are no significant differences between the stability probabilities given by the true and the independent distributions. This suggests that the dependencies between parameters of the system are not directly responsible for the stability of its cell state; rather it is the (feedback) structure of a differentiation cascade imposed by the niche microenvironment that confers stability. Second, there are striking differences between the stability probabilities for the true/independent and the *i.i.d.* distributions (see (Kirk et al., 2015, in press) for further details). In most cases, the *i.i.d.* probability of stability is close to zero: so the structure of the stem cell ecology is far from random. While the structure alone suffices to determine stability, the detailed parameters (e.g. those determining the rates of asymmetric division) will be under the influence of natural selection and will reflect, for example, the physiological requirements for certain numbers/volumes of cells of each given type in a healthy (generally homeostatic) system.

### *Detailed analysis of the stable states of model S*3

More can be learned about the ecology of stem cells and their progeny by investigating the dynamics of these models more closely. In particular it allows us to start to understand the roles of individual parameters on the behaviour of such systems. Here we consider in more detail the possible stable states of model *S*3 (the other models exhibit qualitatively the same behaviour) and investigate what initial states lead to the behaviours associated with each of its biologically relevant fixed points. We find that although two states can be reached that yield positive population sizes for all species, bistability was not observed. This means that there do not exist any experimental conditions within the observed range from which both of the stable states can be reached; rather depending on the system parameters either one or the other will be attained.

We proceed to look at what differences there are in the distributions of parameters leading to each stationary state. From a total sample of 100,000 parameter sets, we find that approximately 8000 lead to fixed point 1 and another approximately 8000 lead to fixed point 2. It is interesting to note that only for this small proportion (16%) of possible parameter combinations states is it possible to reach biologically relevant states; the majority of parameters lead to extinction of species.

In order to ascribe significance to the results we obtain, we need to understand what state each of the fixed points corresponds to. Recall Figure 2 for a graphical depiction of model *S*3: a stem cell gives rise to two progenitor cell populations, which here we call *A* and *B*. Each progenitor cell population can proliferate or differentiate into a corresponding differentiated cell population.

Fixed point 3 corresponds to higher population sizes for progenitor cells by a factor of 10 – for both lineages – compared with the differentiated cell populations. Fixed point 4, in contrast, is characterised by higher levels of differentiated cell populations, again by approximately a factor of 10, relative to the progenitor cell populations. In Figure 3 the distributions of parameters that give rise to these two different states are plotted, along with a description of their meaning.

**Figure 3:**
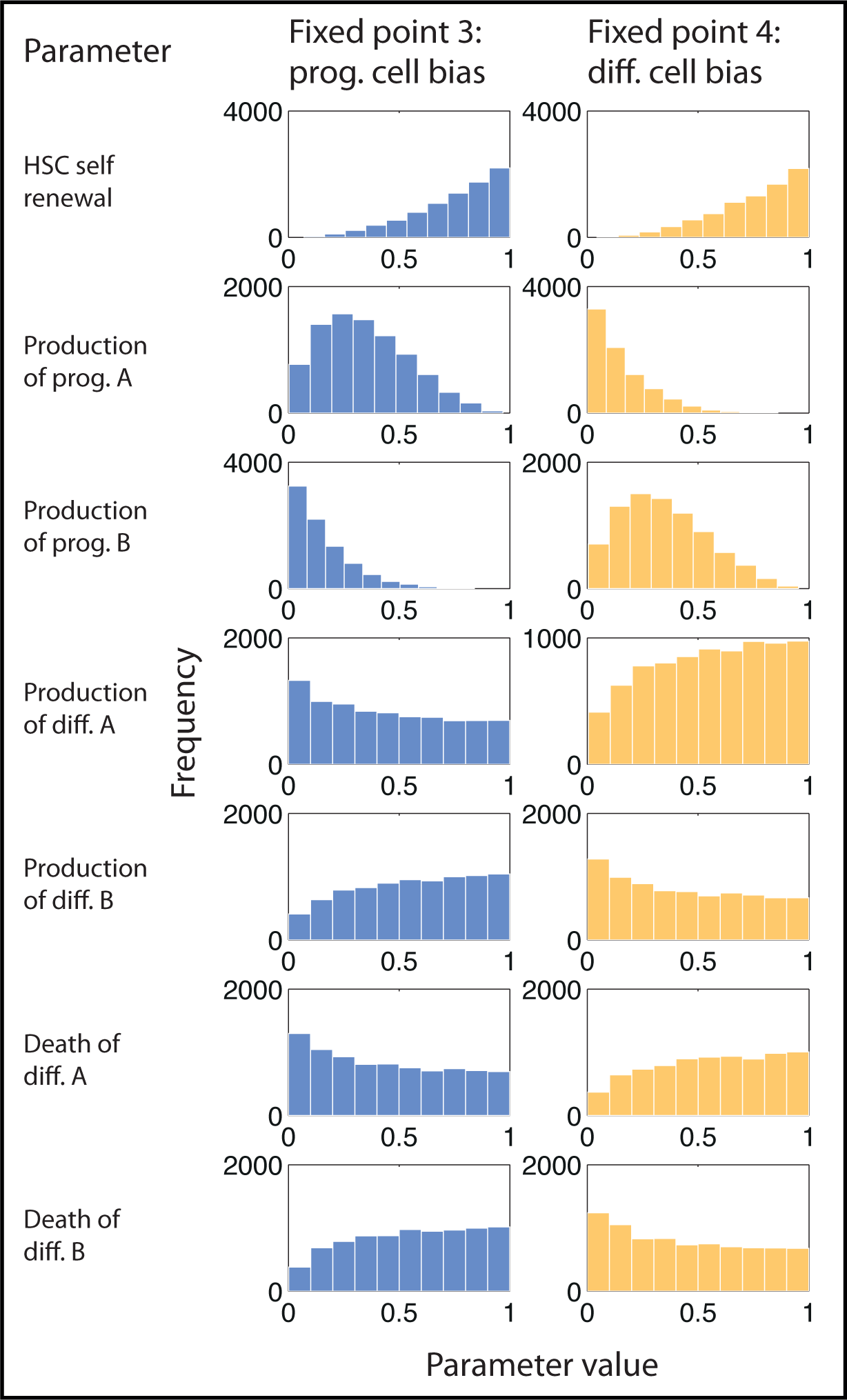
Branching characteristics of distinct lineages. Production rates of progenitor and differentiated cells affect the steady state reached. Histograms of parameter values that lead to one of two fixed points for model *S*3. The first fixed point is characterised by a higher proportion of progenitor cells (prog. cell bias) and the second fixed point is characterised by a higher proportion of differentiated cells (diff. cell bias). ‘A’ and ‘B’ denote the two possible lineages that stem cells give rise to. To reach the progenitor cell-biased state, lineage A is favoured, whereas to reach the differentiated cell-biased state, lineage B is favoured.

By studying Figure 3 we can describe the differences that lead to one fixed point or the other. To reach the state dominated by progenitor cells requires higher production rates of lineage *A* progenitors than lineage *B*. It also requires lower production and death rates of differentiated cells of lineage *A* compared to lineage *B*. We observe symmetries between the distributions that lead to each state: to reach the state dominated by differentiated cells requires, conversely, a higher production rate for progenitor cells of lineage *B* than lineage *A*, and lower production and death rates for the differentiated cells of lineage *B* than lineage *A*.

Describing how lineage bias influences the proportions of different cell species at steady state is especially interesting given the importance of branching fates in stem cell hierarchies, for example in the haematopoietic lineage between myeloid and lymphoid cell fates. The analysis performed here on fixed points 1 and 2 extends the concept of mapping a model’s basin of attraction in parameter space. We can find what regions in parameter space lead to one, or another, fixed point and begin to delineate a boundary between them. Characterising the behaviour of a model with respect to a broad area in parameter space, rather than only at some specific values, enhances our understanding of a model and its potential use.

## Discussion

The ability to make robust fate decisions in a stochastic environment, and the ability to remain a homeostatic population of differentiated, differentiating and stem cells, despite frequently low numbers of stem cells, is a characteristic feature of multi-cellular organisms (Garcia-Ojalvo & Martinez-Arias, 2012; Pauklin & Vallier, 2013; Philpott & Winton, 2014). Such a system can be disrupted, for example, by introducing cells with uncontrolled differentiation and proliferation patterns, i.e. cancer (akin here to an invasive species in classical ecology); then a set of new population dynamics takes over and determines the fate of the cell population. Quite generally, stem cells and their progeny represent populations of interacting cell species, analogous to populations of interacting species in ecology, and are thus amenable to being modelled using concepts from population biology. Substantial research has already been undertaken in ecology and has considered, in particular, the relationship between complexity and stability (Allesina & Tang, 2012; Elton, 1958; MacArthur, 1955; May, 1972; Ives & Carpenter, 2007; Roberts, 1974; Saavedra et al., 2011).

Mathematical analyses allow us to address questions that are inaccessible experimentally. Here we have developed and applied mathematical methods that characterise the stability properties of stem cell population models, focussing on the effects of heterogeneity and of dependencies between species in a hierarchy; while we have increasing experimental power, for example, to do *in vivo* imaging of the haematopoietic stem cell niche in bone marrow (Lo Celso et al., 2009), or to study stem cells in intestinal crypts (Drost et al., 2015), most processes are not directly accessible to observation, and mathematical models can be used to link observables to underlying processes in a rational and hypothesis-driven way. Here the structure of the cell population — stem cells and their progeny — is found to be crucial in enabling stem cell systems (i.e. stem cells, progenitors and their descendants) to reach stable states; homogeneous (randomly interacting) cell populations are not stable. Parametric dependencies affect the stability to a much lesser extent, and we still find stable conditions for stem cells and their progeny to exist in homeostasis when the detailed parameters are ignored.

We analysed the fixed points of one system (model *S*3) in more detail, as we found that multiple cell states could be reached by varying the *in silico* experimental conditions. The balance of progenitor and differentiated cells in model *S*3 is controlled by the propensity of stem cells to favour production of progenitor and differentiated cells in one of two possible lineages. Given the number of possible branching points in adult stem cell hierarchies, characterised by (for example) the multiplicity of haematopoietic progenitor cell species and the possible interactions between them (Wang & Wagers, 2011; Wilson & Trumpp, 2006), such asymmetries are very interesting to identify, and could exert key control over cell fate choice. To analyse this branching process further, comparison of this model to cell species data is required; this will allow us to distinguish between the model’s two lineages.

The number of states and stability properties given by models *S*2 and *S*3 differ, however model *S*4 shares very similar fixed point characteristics to model *S*3. This suggests that whereas niche-mediated feedback onto stem cells is a key factor in state determination, whether the signal comes from the progenitor or the differentiated cell pool (the distinction between models *S*3 and *S*4) is much less important. The fact that structurally different (though related) models share qualitative features is encouraging as this suggests that such feedback might be a generic design feature shared across stem cell systems (Babtie et al., 2014). In light of the results and taken against the background of a vast body of work in population biology, it is certainly hard to propose other mechanisms that would confer such stability.

The definition of stability used throughout this work — that a system at a fixed point will return to the same fixed point following a small perturbation — at times may not match the biologically ‘stable’ properties that we aim to describe. One example of such a mismatch is that of oscillating systems, which can be stable in the sense of persistence. Another example is more subtle: if we compare two bistable systems, one where both fixed points have positive values for all species, and the other where for one fixed point at least one (or more species) goes to 0, we might wish to distinguish between them. That is, we might wish to call the perturbation from one state to another that causes at least one species to vanish greater (in the sense of being destabilising) that the perturbation that causes a state change that is not associated with the extinction of any species. This is an interesting avenue for future work where perhaps different criteria for stability could be used that reflect other aspects of biological homeostasis, such as species’ extinction (Saavedra et al., 2011) or the effects of neutral mutations (Traulsen et al., 2012).

Recent theoretical and experimental studies suggest that multistability plays an important role in cell fate determination, demonstrated via studies of the Wnt signalling pathway (MacLean et al., 2015; Schuijers et al., 2015). While the bistable model of (MacLean et al., 2015) was proved to have two stable states for certain parameter values, similar analysis has not yet to our knowledge been performed for the feedback mechanism proposed in (Schuijers et al., 2015). Generalised stability analysis could shed light on the bistable regime controlled by the Ascl2 gene that is activated by Wnt; a system amenable to modelling. Here, the ecological perspective is perhaps most intimately coupled with cellular and molecular processes, and we can begin to study the multi-level and multi-scale interplay between these different levels *in vitro*, *in vivo* an *in silico*.

## Conclusions

Here we have seen that structure (in the sense of either an underlying interaction network) bestows stability on such systems. We have shown how the stability dramatically decreases when structure is removed. We found this to be the case for all of the stem cell models that we studied here. For models with multiple biological steady states, we identified how each could be reached and in doing so mapped out the basins of attraction in parameter space. This provides insight into how branching decisions in stem cell hierarchies can be made. What emerges from this analysis is the remarkable robustness of stem-cell systems: their stability following a perturbation is a result of cell–cell interactions. This robustness has two consequences: (i) it provides stem cells with a stable ecosystem in which they can fulfill their function in e.g. maintaining tissue homeostasis; (ii) the flip-side is that malfunctioning stem-cell systems — such as systems with additional competition with cancer (stem) cells — may also be robust to a similar extent (MacLean et al., 2015; Youssefpour et al., 2012). Note, however, that the robustness we discuss here in no way limits the ability of stem cells and their progeny to exhibit considerable levels of heterogeneity — this is possible independently of the population dynamics(Rué & Martinez-Arias, 2015). But as our understanding of the structure of stem cell ecosystems (as well as ecosystems more generally) increases we also learn how to shape their fate: being aware that cancer is an evolutionary/ecological disease (Frank, 2007) is now opening up promising new therapeutic interventions.

## Methods

### Model development

Four models are introduced, each consisting of stem (*S*), progenitor (*P*), and differentiated (*D*) cell populations. The first of the models (*S*1) has three cell populations; the remaining three (*S*2 *– S*4) have five. These extra two populations correspond to a branching point in the differentiation hierarchy (for example, in haematopoiesis, into myeloid and lymphoid lineages).

We make the assumptions that (i) renewal is restricted to *S* and *P*; (ii) only *D* are depleted through death/migration; (iii) differentiation is irreversible; (iv) a cell can influence its parent/grandparent population via intercellular signalling. The models are depicted in Figure 2, and full description of their composition including the equations that govern them is given in Appendix A.

### Generalised stability analysis

In order to assess the stability of cell states, we calculate the *Jacobian/community matrix*, for a given state of the system (set of parameter values). This allows us to determine whether or not the system is stable in this state. We repeat this procedure for a large number of parameter sets, sampled in the parameter space in an attempt to capture the global behaviour characteristics of the system. From this analysis, we determine the probability that each fixed point of a given model is stable. We compare these probabilities with those derived from a null distribution, obtained by permuting the connections between cell populations at random. To calculate the independent null distribution we sample with replacement the distribution over each entry in the Jacobian, maintaining the entry position. To calculate the *i.i.d.* distribution we again sample with replacement from the Jacobian, but we now pool entries from all positions, thus the distribution from which we are sampling is now *i.i.d.* (Kirk et al., 2015, in press). Further details of the methods of statistical analysis are given in Appendix B.

## Author’s contributions

ALM, PK and MPHS designed the study and developed the models; ALM and PK analysed the models; ALM, PK and MPHS wrote the manuscript. All authors have approved the final version of the manuscript.

## Acknowledgements

ALM would like to thank Narges Rashidi for helpful scientific and artistic discussions. ALM and MPHS acknowledge funding from the BBSRC through a PhD Studentship; PK and MPHS were funded by HFSP grant GGP0061/2011.

## Appendix A. Mathematical Models of niche-mediated stem cell feedback

We develop four ordinary differential equation (ODE) models that describe stem cell differentiation dynamics. They all share common features and differ in those parts that we wish to compare and contrast. In each model we have a three-layer hierarchy, consisting of stem, progenitor and differentiated cells. Each model is built from a set of shared rules that describe cellular growth, death and differentiation. These rules also define how one species can regulate the growth of another through feedback.

The models differ in the number of species that exert and experience the effects of feedback. Model *S*1 has three species, whereas models *S*2 *– S*4 each have five. The extra two species result from a branching point that represents branching cell fates: when the stem cell divides it is able to differentiate into one of two progenitor species, each of which produces differentiated cell progeny. For example in the haematopoietic hierarchy, the myeloid/lymphoid branching point from a multipotent progenitor represents an early cell fate choice. Further down the myeloid lineage we might reach, for example, the monocyte/granulocyte branching point from a granulocyte monocyte progenitor. All of the *S*-models share a common set of assumptions. Namely, that

- stem and progenitor cells can self renew, but differentiated cells cannot;
- differentiation is irreversible (dedifferentiation is not allowed);
- stem and progenitor species cannot die or migrate out of the niche;
- feedback enters the model through linear growth inhibition of one species on either itself or its parents/grandparents.

The models are parameterised by terms representing growth, differentiation and death rates. We are going to investigate the region of parameter space *p* ∈ [0, 1] for each model parameter *p*. Parameters are assigned biological meaning, thus should never be negative, and we set the upper limit to be 1 so that each *p* is interpreted as a rate parameter. This limit can be set without loss of generality: different parameter ranges could be studied with a rescaling parameter that would not alter the results beyond a rescaling of time (here given in arbitrary units).

### Appendix A.1. *Model S*1

This model describes the dynamics of the following species in the haematopoietic hierarchy: stem cells (*x*_0_); progenitor cells (*x*_1_); and differentiated cells (*x*_2_). Their dynamics are given by

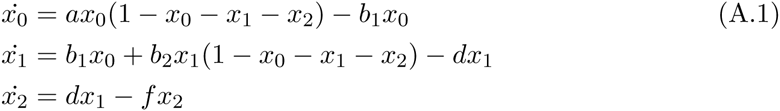

where the parameters (*a, b*_1_, *b*_2_, *d, f*) characterise the phenotypes of the species and are described in Table A.2. Here, populations *x*_0_ and *x*_1_ self-renew at rates *a* and *b*_1_, respectively. They differentiate into daughter species at rates *b*_1_ and *d*, respectively, and *x*_2_ dies/migrates out of the niche at rate *f*. There is also feedback onto the self renewal terms of *x*_0_ and *x*_1_: these are affected by all species in the model.

**Table A.2.**
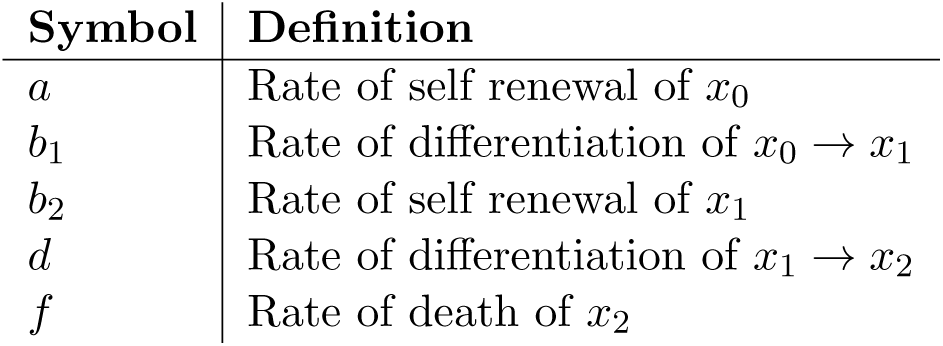
Description of parameters that characterise models *S*1.

| Symbol | Definition |
| --- | --- |
| $a$ | Rate of self renewal of $x_0$ |
| $b_1$ | Rate of differentiation of $x_0 \rightarrow x_1$ |
| $b_2$ | Rate of self renewal of $x_1$ |
| $d$ | Rate of differentiation of $x_1 \rightarrow x_2$ |
| $f$ | Rate of death of $x_2$ |

### Appendix A.2. *Model S*2

Here we include a branching point in the hierarchy and consider feedback from differentiated cells onto progenitor cells, but not onto the parent stem cell population. We now have two progenitor cell populations, *x*_1_ and *x*_2_, that differentiate into two differentiated cell populations, *x*_3_ and *x*_4_, respectively. The equations that specify this model are

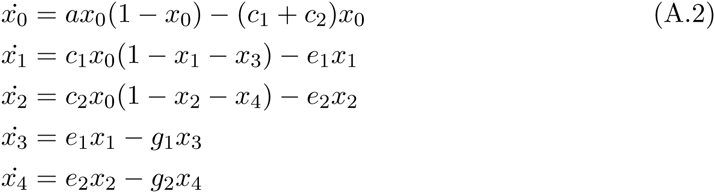

where the parameters (*a, c*_1_, *c*_2_, *e*_1_, *e*_2_, *g*_1_, *g*_2_) are defined in Table A.3.

### Appendix A.3. *Model S*3

This model extends model *S2* by including a separate level of feedback: from differentiated cells onto progenitor cells, in addition to the feedback exhibited by *S2*. The equations for model *S3* are given by

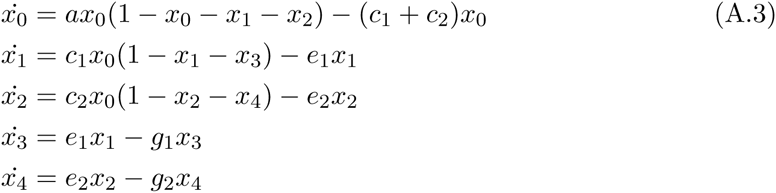

where the same parameter set as for model *S2* describes the model.

### Appendix A.4. *Model S*4

This model considers additional feedback from differentiated cells onto stem cells. So we consider the joint effects of (differentiated *→* progenitor cell) feedback and (progenitor *→* stem cell) feedback. The equations that specify *S8* are

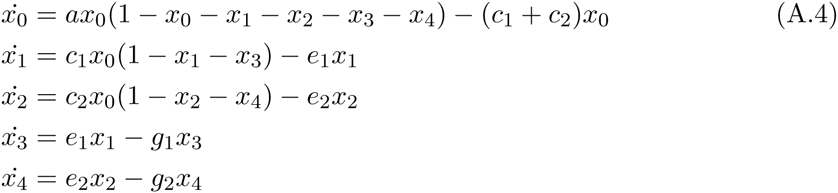

where the parameters are, as above, specified in Table A.3. This completes the family of models that we study here, exhibiting a range of behaviour targeting mechanisms of cell fate decision-making and the role that niche-mediated feedback plays.

**Table A.3.** Description of parameters that characterise models *S*2 – *S*4.

| Symbol | Definition |
| --- | --- |
| $a$ | Rate of self renewal of $x_0$ |
| $c_{1/2}$ | Rate of differentiation of $x_0 \rightarrow x_{1/2}$ |
| $e_1$ | Rate of differentiation of $x_1 \rightarrow x_3$ |
| $e_2$ | Rate of differentiation of $x_2 \rightarrow x_4$ |
| $g_1$ | Rate of death of $x_3$ |
| $g_2$ | Rate of death of $x_4$ |
| $p_{1/2}$ | Rate of self renewal of $x_{1/2}$ |

## Appendix B. Extended methods for linear stability analysis

Linear stability analysis describes a method to assess how a dynamical system responds to small perturbations about its equilibria. Here we introduce linear stability analysis and describe how we use generalised stability analysis to study the stability properties of fixed points (Kirk et al., 2015, in press).

### Appendix B.1. Background

We begin with some definitions. A *system* is a set of interacting dynamic variables. Since here we are considering systems such as those found in an ecological setting, we refer to the variables as *species*. We consider (eco)systems whose dynamics can be described by ODEs, so we can write down the system dynamics in general form as

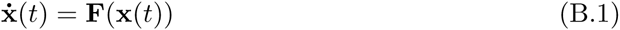

where **x**(*t*) is a vector of size *n* (*n* is the total number of species) that gives the abundance of each species at time *t* (we will use **x**(*t*) = **x** i.e. time dependence implicit). The abundance could be represented by whole numbers of species or by their concentration, given by the fraction of a species occupying a certain space. **F**(**x**(*t*)) = [*F*_1_(**x**(*t*)), *F*_2_(**x**(*t*)), …, *F_n_*(**x**(*t*))] are non-linear functions that define the rates of change of the species and include the model’s parameter dependencies.

Since we are interested in the behaviour of the model close to steady state, we consider a Taylor expansion of *F* (**x**(*t*)) about the point **x_0_**, where **x_0_** defines a fixed point of the system. In doing so we aim to learn about how the model responds to perturbations from this steady state. Discarding terms of order *x*^2^ or higher from a Taylor expansion we have

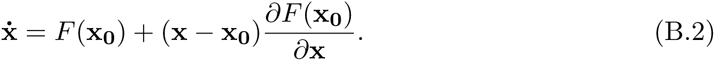

Since **x_0_** is a fixed point of the system, *F* (**x_0_**) = 0. Substituting **y** = **x** – **x_0_** we have the linearised system

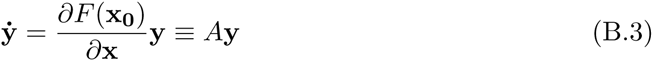

where *A* is called the Jacobian matrix in a mathematical context or the community matrix in an ecological one (Levins, 1968). We can study *A* to determine properties of the system at hand. Specifically, we can find the eigenvalues of *A* (λ_*i*_ where *i* ∈ (1, 2, …, *n*)) by solving

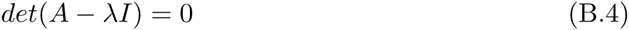

for λ, where *I* is the *n*×*n* identity matrix. We can then use the *λ* to define stability criteria: if *Re*{λ_*i*_} < 0 ∀*i* then the system is *stable* about the point **x_0_**: if the system is perturbed slightly from state **x_0_**, it will return to the same state following some time delay. If this condition does not hold, the fixed point is unstable. See, for example, Strogatz (1994) for further discussion of linear stability analysis. This definition of stability is used in a broad variety of applications in dynamical systems; we are interested particularly in its ecological applications, by May (1973) and others.

In 1972, Robert May proved a theorem that demonstrated that as the complexity of a system increases, its stability will in general decrease and for large enough systems tends towards zero (May, 1972). Complexity here is quantified either by the total number of species in the system or by the number of links between them. For a randomly connected system of *n* species, with connectance — the number of actual links over the number of all possible links — *C* and average link strength (or variance) *α*, then, for large *n*, a system will almost certainly be stable when

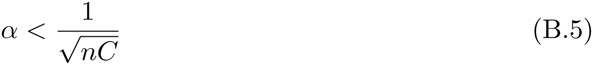

(May, 1972). This result contradicted current opinion of the time which stated that in general as systems grow in complexity their stability should increase (Elton, 1958; MacArthur, 1955). The implications and uses of this theorem have been wide and varied; outside of ecology it has found application in fields including molecular biology and economics.

Given this theorem, a question arises, namely, how do large systems persist in nature? As their size and complexity increases, Equation B.5 states that their probability of stability will eventually approach zero. To begin to answer this, note that Equation B.5 gives a statement about the behaviour of systems where links between species are placed at random. Of course in the real world interactions are not completely random but guided by evolution.

**Figure B.4:**
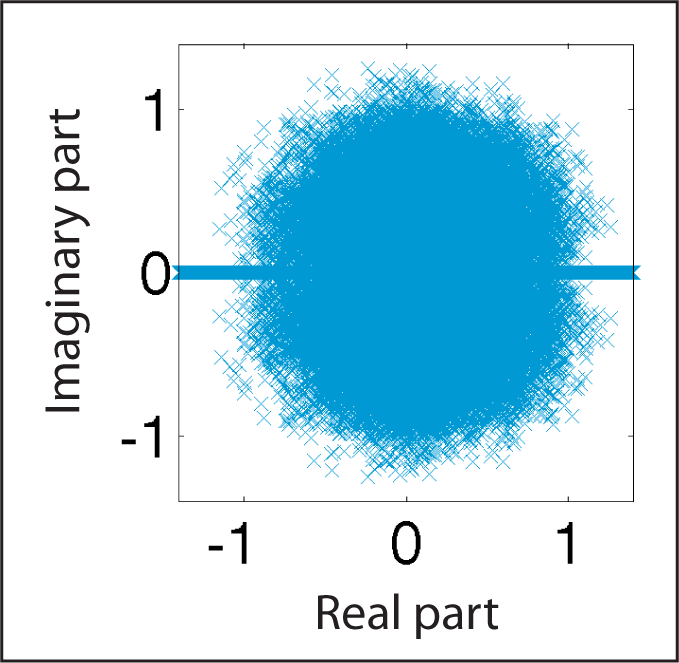
*Illustration of the circular law for a random matrix distribution.* Jacobian entries for 5 *×* 5 matrices are drawn from an empirical *i.i.d.* distribution. The eigenvalues for each Jacobian are calculated and can be plotted on the complex plane, as shown here.

Results regarding the stability of biological systems by May, and recent extensions to his work by Allesina & Tang (2012), rely on interactions between species being *i.i.d.* because they appeal to a theorem known as the circular law, and its extensions (Girko, 1984; Tao et al., 2010). This states that, given a system of *n* species with interactions between them drawn according to an *i.i.d.* distribution, then in the limit *n* → ∞, the eigenvalues of the system lie on a circle in the complex plane. In the case of finite *n*, the spectral distribution of eigenvalues still approaches a circular distribution, with some “bleeding” at the edges, especially along the real axis. As an example, shown in Figure B.4 are the eigenvalues of 100,000 5 × 5 matrices with *i.i.d.* entries. Illustration of the circular law is useful as a point of comparison with numerical results.

### Appendix B.2. Generalised stability analysis

In order to conduct stability analyses for each of the models we evaluate the Jacobian matrix at each fixed point for a given parameter vector. We sample the parameter space randomly, keeping only those points that give rise to fixed points in the positive quadrant, we discard parameters giving rise to fixed points that are not feasible in the sense of being biologically relevant or *reachable*, from some initial conditions.

We follow these steps to determine the stability probability for a given model around a fixed point:

1. Solve 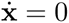 to find symbolic expressions for the values of each species (total number *n*) at each fixed point in terms of the parameters.
2. Sample a 1 *×* d parameter vector **p** (model contains *d* free parameters) and evaluate the fixed point for these values. Check *reachability*: if the fixed point lies within the positive orthant, i.e. all species ≥ 0, proceed. If not, resample.
3. For **p**, find the eigenvalues λ_*i*_, *i* ∈ (1, 2, …, *n*) of the Jacobian, *J*. Recall that 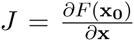 where *F* is given by the model equations, 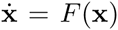 and evaluated at fixed point **x_0_**.
4. Repeat steps (2) and (3) until a total of *N* reachable points are found.
5. Construct **s**, where *s*_*j*_ = 1 if *max*(*Re*{λ_*i*_}) < 0 for λ_*i*_ of *p*_*j*_ ∈ **p**, and *s*_*j*_ = 0 otherwise. i.e. *s*_*j*_ indicates whether or not the model is stable around the fixed point for *p*_*j*_.
6. Calculate the stability probability: 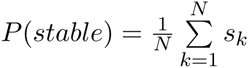

**Figure B.5.**
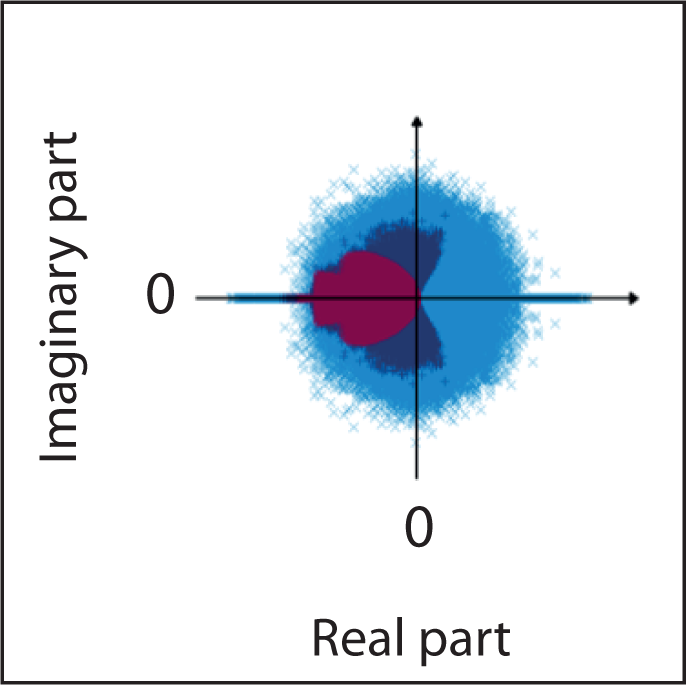
Eigenvalue distributions for the true, independent and *i.i.d.* distributions of Model *S*4. The independent distribution (dark blue) deviates considerably from the near-circular *i.i.d.* distribution (light blue); the true distribution (purple) is even more tightly constrained.

Note that the stability probability *P* (*stable*) = *P* (*stable*|*reachable*). That is, we calculate the probability that a given fixed point is stable given that it is reachable from the starting point (parameter values and initial conditions).

Aside from calculating *P* (*stable*), given the distribution of Jacobian matrices that this analysis gives us, we can also investigate the fixed points of a model in other ways, such as by quantifying the basins of attraction of a model and looking at their intersection. We can also look at basins of attraction in parameter space: the effects that varying parameters (rather than initial conditions) have on the reachability of the fixed points.

### Appendix B.3. Null distributions to describe the stability of a model

In this work we set out to study the stability properties of stem cell models and in particular to determine which properties of the models confer to them more or less stability around a certain fixed point. So, in addition to comparison between models, for a single model we would like to compare its stability properties with those of a permuted version of the same model. In doing so we essentially create a null distribution for the stability properties of a model around a given fixed point.

In fact, we consider two null distributions for each model. First, we sample with replacement from the empirical distributions that we have for each entry in the model Jacobian, maintaining the entry place in the matrix – denote this distribution *ϕ*_*i,j*_. For example, to sample from *ϕ*_1,1_ is to sample from the *N* values we have for the first entry in the Jacobian: *J*_1,1_. This we call the independent null distribution: it removes dependencies between entries in the Jacobian. Second, we sample again from the distribution over all Jacobian entries but now we combine the values for all entries so that, for each entry in our new Jacobian, we sample with replacement from the empirical distribution containing the *n*×*n*×*N* values – we denote this distribution *θ*. This is an empirical *i.i.d.* distribution since we are now sampling independently from the same distribution for each entry in the Jacobian.

To construct these two null distributions we follow steps (1)-(6) as above but replace (3) by, for the independent null distribution:

- Construct a new Jacobian by sampling with replacement from *ϕ*(*i, j*) for each entry *J* (*i, j*) and find its eigenvalues

and for the *i.i.d.* null distribution:

- Construct a new Jacobian by sampling with replacement from *θ* for each entry in *J* and find its eigenvalues.

Given these distributions, which define independent and *i.i.d.* permutations of the original eigenvalue distribution of a model, we can investigate the impact that the structure of a model has on the eigenvalue distribution. Shown in Figure B.5 are the true, independent and *i.i.d.* eigenvalue distributions for fixed point 4 of model *S*4. Here we see that the eigenvalues of both the true and independent distributions are more tightly constrained in the complex plane than the eigenvalues of the *i.i.d.* distribution, and that the *i.i.d.* distribution approaches a circular distribution conforming with the circular law (Girko, 1984), and in similarity with Figure B.4.

